# A migration-associated supergene reveals loss of biocomplexity in Atlantic cod

**DOI:** 10.1101/361691

**Authors:** Tony Kess, Paul Bentzen, Sarah J. Lehnert, Emma V.A. Sylvester, Sigbjørn Lien, Matthew P. Kent, Marion Sinclair-Waters, Corey Morris, Paul Regular, Robert Fairweather, Ian R. Bradbury

## Abstract

Intraspecific phenotypic diversity is integral to ecological resilience and the provision of ecosystem services^1^. Chromosome structural variation may underpin intraspecific diversity and complex phenotypes^2^ by reducing recombination within supergenes containing linked, co-adapted alleles. Connecting ecologically-relevant phenotypes to genomic variation can enable more precise conservation of exploited marine species by protecting important genetic diversity^3,4^. Here, using genome-wide association analysis of a 12K single nucleotide polymorphism (SNP) array we confirm that an ancient, derived chromosomal rearrangement consisting of two adjacent inversions is strongly associated with migratory phenotype and individual-level genetic structure in Atlantic cod (*Gadus morhua*) across the Northwest Atlantic. The presence of all identified migration-associated loci within this rearrangement indicates that pervasive variation in migration phenotype is in part controlled by a recombination-resistant supergene, facilitating fine-scale individual phenotypic variation within Northern cod. Furthermore, we reconstruct trends in effective population size over the last century, and find genomic signatures of population collapse, and different patterns of population expansion and decline among individuals based on supergene alleles. We demonstrate declines in effective population size consistent with the onset of industrialized harvest (post 1950) and substantially reduced effective size of individuals homozygous for the derived chromosomal rearrangement relative to heterozygous individuals or those homozygous for the ancestral version of this chromosomal region. These results illustrate how chromosomal structural diversity can mediate fine-scale genetic and phenotypic variation in a highly connected marine species, and suggest a loss of biocomplexity from a migration-associated supergene within Northern cod by overfishing.

---

Consideration of intraspecific variation is central to the effective management of natural resources and ecosystem services^5, 6^. Individual phenotypic and genetic variation can play a key role in dictating ecological composition and function, and accordingly perturbations to this variation have been shown to alter ecosystem structure^7^. Incorporating intraspecific diversity into management plans requires the identification of heritable phenotypes linked to ecological resilience and biocomplexity^1, 8^. Genomic analyses have increasingly revealed chromosome structural rearrangements (e.g., inversions, translocations) underpinning ecologically variable traits, supporting an important role for genomic architecture in promoting intraspecific ecological diversity^9, 10^. For example, chromosomal inversions may facilitate the evolution of complex phenotypes in sympatry through the formation of supergenes or clusters of linked adaptive loci in regions of reduced recombination^9^. Quantifying the relationship between genomic architectural variation and ecologically relevant phenotypes can provide deeper understanding of the genomic drivers of biocomplexity, enabling genomics-guided species management and precise measurement of human impacts within species^11^. However, the relationship between genomic architecture and key ecological traits remains largely unknown in most exploited species ^4^

Atlantic cod (*Gadus morhua*) have an extensive history of exploitation and increasing evidence of genomic structural variation associated with ecological adaptation^4, 10, 12^. The Northern cod population in northwest Atlantic waters around Newfoundland and Labrador have undergone multiple population declines, the most recent and drastic of which has been driven largely by overharvesting following adoption of industrial-scale fishing, in tandem with climate shifts^13^. In conjunction with recent declines in abundance, size and age at maturity have also declined within the Northern cod stock since the 1970’s^14,15^. Both the presence of a genetic basis for intraspecific variation in the species, and reports of changes in key life history traits suggests the potential that fishery-induced selection has occurred^15^.

Individual and regional genomic diversity that differentiates Atlantic cod populations by environment and migratory behavior has been identified across the species range in the North Atlantic^12^. Recent genomic analyses have revealed large chromosomal rearrangements on four linkage groups (LG1, LG2, LG7 and LG12)^10,12^. Two adjacent inversions within LG1^16^ and the rearrangement on LG12 differentiate offshore migratory and coastal non-migratory populations^4, 10, 16, 17^. The migratory phenotype exhibited by Northern cod may have imposed increased vulnerability to overfishing in these populations. However, the link between these rearrangements, migratory phenotype, and the consequence of overharvest remains largely unexamined. Understanding the link between phenotypic variation, population stability, and genomic architectural variation is particularly important in Northern cod, which have undergone one of the most severe population crashes recorded in an exploited marine species, and have failed to recover to historical abundance despite a lengthy and continuing fishing moratorium^19^.

Here we explore the genomic basis of individual variation within the Northern cod stock around Newfoundland and Labrador (Figure 1A, Supplementary Table 1) using a 12K single nucleotide polymorphism (SNP) array. We examine the role of chromosomal rearrangements associated with migration phenotype and temporal trends in effective population size over the last century. Within the Northern cod stock, principal component (PC) analysis using 6669 informative SNPs identified three distinct clusters along the first PC axis (Figure 1B). Outlier identification using the Mahalanobis distance for each SNP and the first PC revealed individual genetic structuring was greatest among SNPs localized between 9.89 and 27.22 Mbp on LG1, consistent with boundaries of the derived chromosomal rearrangement consisting of two adjacent inversions on LG1^16^. These inversions arose in the Eastern Atlantic 1.6 – 2.0 Mya, prior to the most recent colonization of the Northwest Atlantic^10,16^, suggesting trans-Atlantic introduction of the derived LG1 rearrangement by migratory cod. The positions of outlier loci indicate observed clusters correspond to individuals homozygous for the ancestral collinear chromosome orientation, and those homozygous or heterozygous for the derived LG1 rearrangement haplotype (Figure 1C). The presence of two adjacent inversions may prevent multiple crossovers events, suppressing recombination along the entire rearrangement and enabling this genomic region to function as a single supergene^10, 20^.

**Figure 1.**
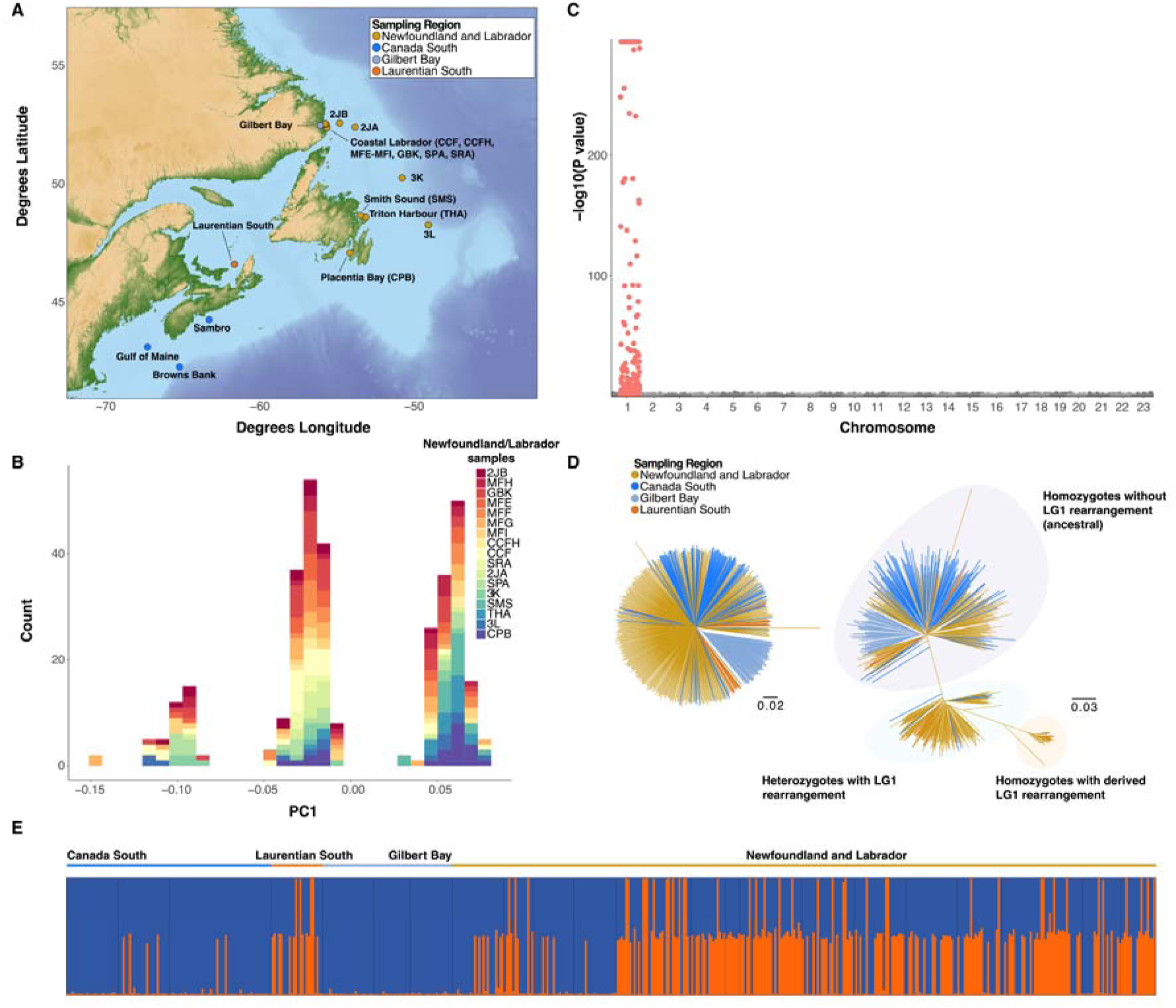
Map of Atlantic cod (*Gadus morhua*) sampling locations in the northwest Atlantic and genetic structuring across all sampled sites and within Northern cod. **A)** Sampling sites across North America colour-coded by population **B)** Genetic structure within Northern cod visualized using the first principal component (PC) calculated from 6030 SNPs in *pcadapt.* Colours correspond to sample groups taken from sites within the greater Northern cod population **C)** Genomic sources of individual level-genetic variation Northern cod assessed by significance of correlation (*p* value) of each SNP locus with the first PC axis obtained from transformed Mahalanobis distances. SNPs within the LG1 rearrangement are coloured in red. **D)** Individual neighbour joining trees for 511 individuals from all sampling locations calculated using 2500 randomly selected SNPs outside of Northern cod inversions (left) and 237 SNPs in the LG1 rearranged region(right) **E)** Bayesian clustering results from STRUCTURE for 237 SNPs in the LG1 rearranged region calculated among all 511 individuals from all sampling locations for K=2 genetic clusters.

Population structure analysis was expanded geographically to include samples from throughout the Northwest Atlantic in waters around Newfoundland and Labrador, Southern Canada, the Laurentian Gulf, and coastal Labrador. We conducted separate analyses using genome-wide SNPs from regions without known inversions (n=2500), and SNPs within the rearranged region on LG1 (n=237). Individual Neighbour-Joining trees revealed little structuring at neutral loci, and again uncovered the presence of three discrete groups using only SNPs within the LG1 rearranged region (Figure 1D). Bayesian clustering analysis of SNPs within the LG1 rearranged region also demonstrated ancestry proportions consistent with homozygous or heterozygous genotypes from ancestral or derived rearranged LG1 haplotypes (Figure 1E).

To quantify the relationship between chromosomal rearrangements in the Atlantic cod genome and migratory phenotype, we conducted genome-wide association analyses across all sampled populations. We assigned regional-level migratory and non-migratory phenotypes to individuals from each sampled location using previously collected migratory behaviour data^21^ (Supplementary Table 1), and detected migration-associated SNPs using random forest, partial redundancy analysis, and latent factor mixed models. We uncovered 21 SNPs (Supplementary Table 2) associated with migratory phenotype across all three association methods, all located within 9.89 and 27.22 Mbp in LG1, corresponding to the position of the rearrangement (Figure 2A, B, Supplementary Figure 1), and exhibiting high linkage disequilibrium (r^2^ = 0.82, sd = 0.09). Additionally, we detect evidence of a selective sweep within the derived LG1 rearrangement (Figure 2C). A large suite of functional variants (>300) differentiated between coastal and migratory ecotypes in Norwegian waters have previously been identified within this rearrangement^16^. Several of these alleles are suggested to affect swim bladder function and muscular efficiency, facilitating vertical movement at variable depths in offshore sites and enhancing muscular capacity for strenuous migration^10, 16^.

**Figure 2.**
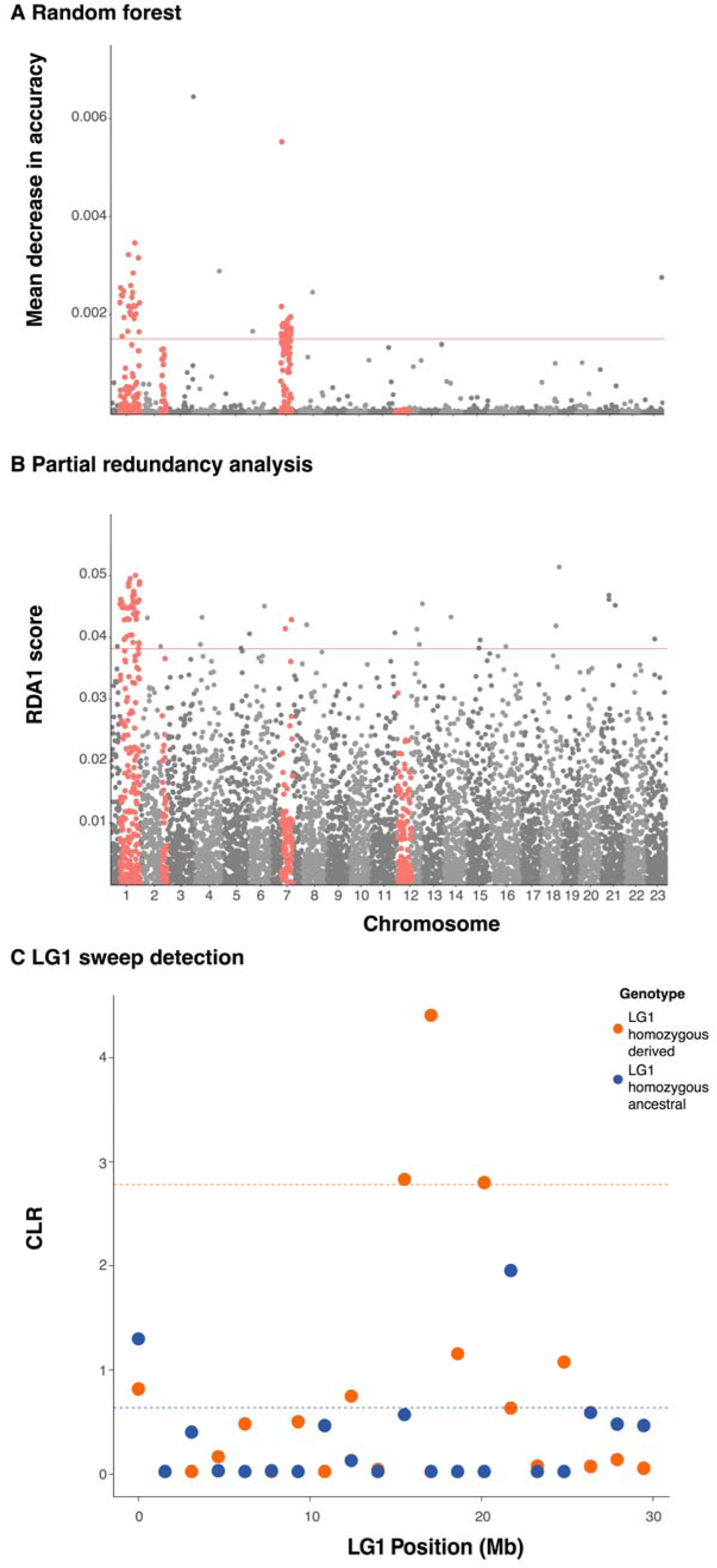
Genome-wide association of 6669 SNPs with migration phenotype assigned by Robichaud and Rose^21^, across all sampling locations, and selective sweeps within LG1. **A)** Mean decrease in accuracy per SNP in Random Forest prediction of migratory phenotype, categorizing SNPs with mean decrease in accuracy > 0.015 as significantly associated. SNPs within rearrangements are coloured in red. **B)** Absolute values of scores for each SNP along the first redundancy canonical axis (RDA1) in partial redundancy analysis of migratory phenotype conditioned by geographic location, treating all SNPs in the 99^th^ percentile as significantly associated **C)** Selective sweeps in LG1 homozygous ancestral and LG1 homozygous derived individuals detected by calculating composite likelihood ratio (CLR) per 20 SNP window with regions exceeding the 90^th^ percentile considered likely candidates for selection

Northern cod experienced substantial overharvest, reflected in the occurrence of one of the biggest population crashes of any marine vertebrate that resulted in >90% decline in abundance during the 20^th^ century^14, 18, 22^. Despite the scale of stock collapse, previous neutral molecular marker-based attempts have failed to detect genetic signatures of this decline^23^. Here, we find that genomic data can detect declines specific to groups with ancestral or derived copies of LG1. We separated individuals into LG1 homozygous derived, LG1 homozygous ancestral and heterozygous groups and then estimated trends in recent effective population size (Ne) in each group using genome-wide SNPs outside of LG1 and LG12 rearrangements identified in Northern cod. We then compared proportional change in Ne relative to maximum observed effective size within each group during the past 150 years to observed and reconstructed stock abundance^13^. We found a similar starting effective population size at the 1860 time point for each group (Ne = 1976 – 2629, Supplementary Table 3), but observed different patterns of expansion and decline between LG1 homozygous ancestral, LG1 homozygous derived, and heterozygous groups (Figure 3A, B), suggesting different selective histories. Within LG1 homozygous ancestral and heterozygous groups, we observed an historical expansion of effective size, indicated by large Ne estimates with confidence intervals including negative values (i.e. infinite), coinciding with a period of population productivity from 1900 to approximately 1970^13^. From 1970 to present, we observe a shared pattern of decline but different contemporary effective sizes of each group. Although non-independence of SNPs may reduce confidence in exact estimates of Ne^24^, the magnitude of different contemporary Ne values between LG1 homozygous ancestral (Ne = 9697) and derived (Ne = 165) individuals indicates potential for loss of migratory phenotype due to overharvest. Comparing LG1 rearrangement groups by region revealed decline in LG1 homozygous ancestral and heterozygous individuals driven by decline in sites around Labrador, suggesting spatial variation in selection intensity (Supplementary Figure 2). The capacity of genomic data to detect recent collapse of Northern cod was also supported by allele frequency-based bottleneck tests.

**Figure 3.**
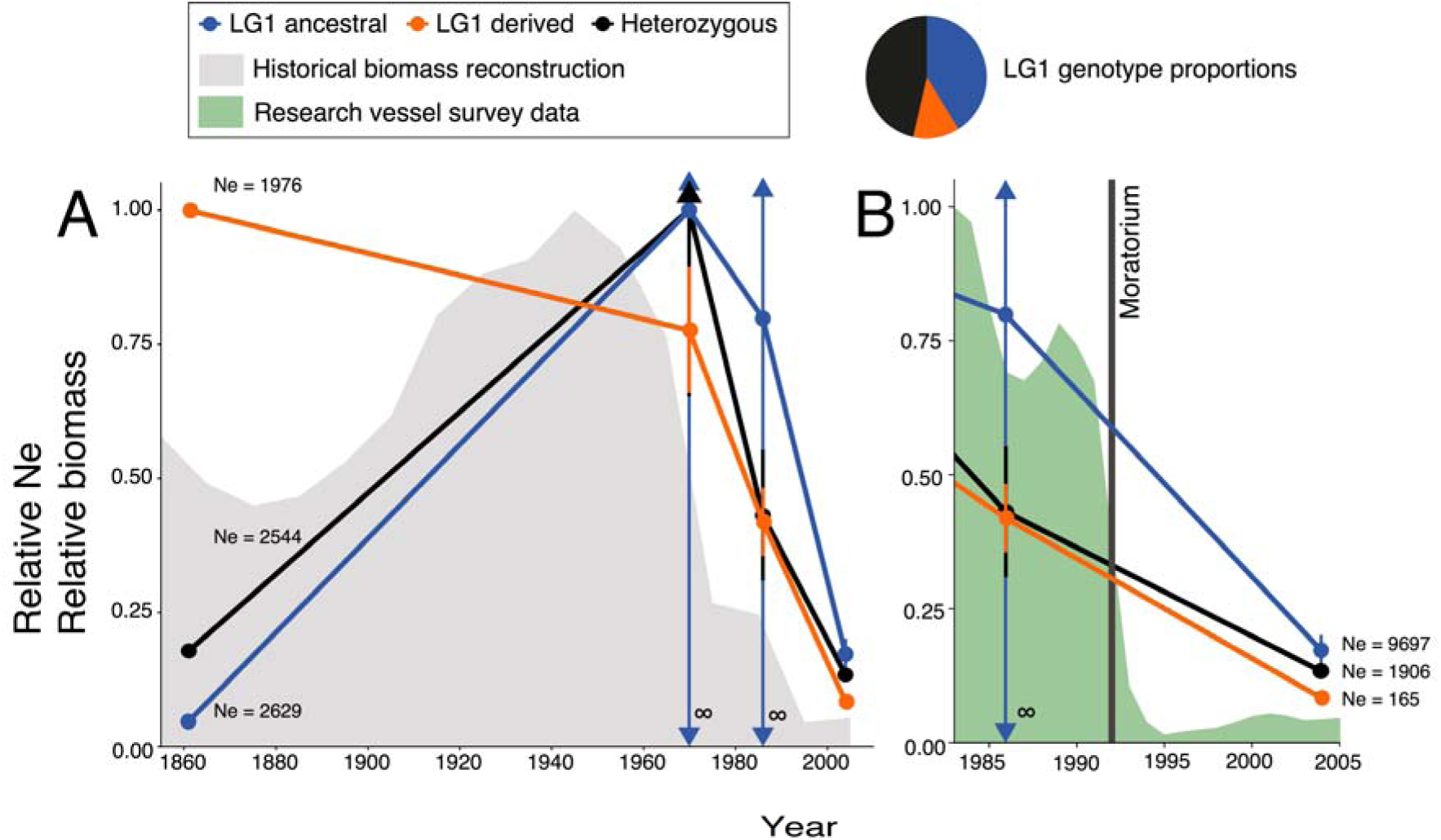
Comparison of recent relative effective population size (Ne) in LG1 homozygous ancestral, LG1 homozygous derived and heterozygous individuals and measured and historically reconstructed stock abundance of Northern cod. **A)** Recent effective population size relative to maximum effective population size within each group of LG1 homozygous ancestral, LG1 homozygous derived and heterozygous Northern cod groupings calculated using LinkNe, plotted with reconstructed estimates of historical biomass measured from 1850 – present^12^ relative to maximum value. Confidence intervals with arrows indicate very large Ne. **B)** Recent proportional effective population size relative to maximum effective population size of LG1 homozygous ancestral, homozygous LG derived and heterozygous Northern cod calculated using LinkNe compared to measurement of stock abundance from 1983 – 2004 relative to maximum value.

Our results provide novel advances in understanding biocomplexity at a genomic level in marine species. We demonstrate that intraspecific diversity in migratory behavior is associated with a derived chromosomal rearrangement consisting of two adjacent inversions spanning LG1 and illustrate that this rearrangement enables extensive differentiation among individuals within Northern cod. Suppressed recombination between opposite orientations of the LG1 rearrangement can promote coordinated inheritance of migration-associated loci, allowing the two adjacent inversions to function as a single supergene protected from gene flow^2, 9, 10, 20^. This supergene is associated with genetic signatures of extensive decline in homozygous individuals, potentially driven by overharvesting. Collapse of Northern cod has resulted in massive restructuring of coastal shelf ecosystems^25^. Continued overharvesting of the derived, migration-associated LG1 rearrangement could lead to widespread altered ecological function and composition in the Northwest Atlantic through changing cod distribution and site persistence. Removal of this genomic diversity through overfishing may also reduce the buffering effect provided by phenotypic diversity within these populations, increasing risk of future collapse^1^. Management of Northern cod stocks may thus require consideration of the genomic architectural diversity within these populations to ensure maintenance of biocomplexity across the Northwest Atlantic.

## Methods

### Sampling and Genotyping

We combined Atlantic cod (*Gadus morhua*) genotype data from three studies conducted by Sinclair-Waters *et al.* ^10,17^ and Berg *et al*.^12^ (Supplementary Table 1). The combined dataset comprised 511 individuals in 24 sample groups collected between 2001 and 2015 from across 18 sites the Northwestern Atlantic. Sampling site details, sample collection and preparation methods are described in Sinclair-Waters *et al*.^10,17^ and Berg *et al*.^12^. All sampled individuals were genotyped using a 12K SNP array developed for Atlantic cod at the Centre for Integrative Genetics (CIGENE), Norwegian University of Life Sciences in Ås, Norway^26^. Following correction for strand flips, we retained a subset of 6669 polymorphic SNPs with complete mapping information across all datasets for further analysis.

### Genetic structure detection

To characterize genomic sources of individual variation within Northern cod, we used the *pcadapt*^27^ R package to conduct principal component analysis (PCA) on the 330 individuals from sampling sites around Newfoundland and Labrador. Using 6030 SNPs with minor allele frequency greater than 0.025, we carried out PCA retaining the first PC axis to identify the largest source of individual genomic variation. We plotted*p* values from transformed Mahalanobis distances measuring significance of correlation with each SNP and this PC axis across each linkage group using the R package *ggman*^28^.

### Range-wide phylogeny and genetic structure analysis

To identify intraspecific variation in LG1 rearrangement frequency relative to the genome-wide average across the northwestern Atlantic, we then calculated individual neighbor-joining trees using all 511 samples. We calculated separate trees for the 237 SNPs with map coordinates within the LG1 rearrangement and for 2500 randomly selected SNPs outside of the known chromosomal rearrangements found in Atlantic cod on linkage groups 1, 2, 7, and 12^12^. Trees were calculated using POPULATIONS v1.2.33^29^, based on Cavalli-Sforza and Edwards^30^ chord distances with 1,000 bootstrap replicates on loci. Neighbour-joining trees were visualized using FigTree v1.74^31^. Subsampling of genome-wide and inversion loci was conducted using the genepopedit R package^32^.

Next, we conducted Bayesian clustering analysis of all 511 sampled individuals with three replicate Markov-Chain Monte Carlo runs on LG1 rearrangement genotypes in STRUCTURE v2.3.4^33^. We implemented these runs using *parallelstructure^34^* in R, assuming K=2 clusters, and conducting 100,000 burn-ins and 500,000 iterations. We visualized individual genetic ancestry proportions using CLUMPAK^35^. Conversion of genotypes to STRUCTURE format was carried out using PGDSpider^36^.

### Genome-wide association analysis of migratory phenotype

We assigned migratory behavioural phenotypes to all individuals in each sampling site from previously identified behavioural classes categorized in Atlantic cod by Robichaud and Rose^21^(Supplementary Table 1). When multiple locations within the range of a sampling site exhibited variable phenotypes, we assigned the most frequent phenotype identified across these locations. Due to potential variation in sampling and behavioural measurement methods between studies, we binned three migratory (M) behavioural classes together (accurate homers, AH, inaccurate homers, IH, and dispersers, D) as migratory (M), and categorized sedentary (S) individuals as non-migratory (NM). We then conducted genome-wide association using two polygenic methods, random forest (RF)^37^ and partial redundancy analysis (RDA)^38^ and a single locus method using latent factor mixed models (LFMM)^39^.

To find polygenic associations with migratory phenotype, we first used random forest^37^ classification, a non-parametric, decision tree-based algorithm. We produced a matrix with individual migratory phenotypes as the response variable, and SNP genotypes at each locus as predictor variables. Random forest was run using the R package *randomForest*^40^ after imputing missing genotypes using the *rfImpute* function. We ran 5000 trees as predictor rank stabilized was ensured with this quantity of trees. We set the mtry parameter to default following testing of optimal mtry parameter values using the *tuneRF* function. Stratified sampling was applied to ensure unbiased representation of migratory phenotype class within each tree. Out-of-bag mean decrease in accuracy was calculated across all trees for each SNP as an estimate of predictor importance. We retained all SNPs with mean decrease in accuracy greater than 0.0015; values below this limit exhibited a sharp drop in mean decrease in accuracy, indicating relatively limited utility of these SNPs in classifying migratory and non-migratory individuals.

We then carried out a constrained ordination using partial redundancy analysis. This method maximizes the proportion of variance explained in linear combinations of response variables by linear combinations of predictor variables^41^. To detect polygenic associations, we used a matrix of diploid genotypes for each individual at each locus as dependent variables, and migratory phenotype as the predictor variable and carried out redundancy analysis using the *vegan*^42^ R package. Missing genotype calls were imputed for each individual using LinkImpute^43^. To account for population structure due to geography, we conditioned the relationship between genotype and migratory phenotype by latitude and longitude. Significance of genotype-phenotype association was calculated with 999 permutations (p < 0.001). We categorized SNPs with absolute values of canonical redundancy axis scores exceeding the 99% percentile on the first axis (RDA1) as significantly associated with migratory phenotype. Linkage disequilibrium was calculated between migration-associated SNPs identified across all three methods in PLINK v 1.90^44^ using an r^2^ cut-off of 0.2.

Detection of SNP association in LFMM was conducted by using a matrix factorization algorithm to calculate loadings of K latent factors, similar to a reduced set of principal components, that each describe a different source of population structure^39^. We empirically estimated the value of K with sparse nonnegative matrix factorization^45^ using the *sNMF* function on all 6669 SNPs in the *LEA* R package^46^. We then used the *lfmm* function in *LEA* to identify SNP association with migratory phenotype using K=4 latent factors. We converted p values for all loci to *q* values to control for false discovery rate^47^ and retained loci with *q* < 0.05.

### Detection of selective sweeps

To identify genomic regions exhibiting signatures of selection on migratory phenotype within Northern cod, we calculated composite likelihood ratio^48^ (CLR) across LG1 using the SweeD^49^ software package. We separated individuals from Newfoundland and Labrador sampling sites into LG1 homozygous ancestral and LG1 homozygous derived, rearranged classes based on individual PC scores. We then calculated CLR for windows of 20 SNPs within derived (n = 41) and ancestral (n = 136) individuals, assuming a folded site-frequency spectrum, and selected regions exhibiting CLR exceeding the 90% percentile as potential targets of selection.

### Contemporary and historical effective population size estimation

To detect the recent genetic history of Northern cod, we used the linkage disequilibrium (LD) based method implemented in LinkNe^50^ to estimate effective population size (Ne). We calculated Ne across a 150 year period in the 330 Northern cod samples from Newfoundland and Labrador. We separated LG1 homozygous derived, heterozygous (n = 153) and LG1 homozygous ancestral individuals based on PC score. Prior to Ne estimation, we filtered genome-wide SNP datasets for these individuals to remove SNPs within the LG1 and LG12 rearrangements identified within Northern cod, resulting in a final dataset of 6270 SNPs. These regions were removed as selection and reduced recombination within rearrangements regions may skew LD inference of historical Ne. For each group, we calculated Ne separately in LinkNe in bins of 0.05 Morgans for all SNPs with minor allele frequency values greater than 0.05. We binned Ne estimates into generations calculated from recombination rate, assuming a mean generation time of 6 years^15^. We assigned a contemporary sampling date of 2010, corresponding to an intermediate estimate of sampling dates from all Northern cod samples. To compare changes in effective population size between groups that exhibited large differences in Ne, we calculated relative Ne by scaling Ne values within each group by the maximum observed Ne for that group. We then plotted relative Ne estimates as proportions against estimates of cod abundance measured from 1983 to 2004, and historical reconstructions of stock biomass from 1850 to present^12^.

To account for the possibility of observed decline in LG1 homozygous derived individuals driven by small sample size, we generated 10 subsamples of 41 LG1 homozygous ancestral individuals and conducted replicate effective size estimation. We observed similar starting effective sizes across replicates (mean Ne = 2577.449, sd = 170.1235) and identified expansions coinciding with stock recovery in all 10 samples. We did not observe patterns of decline matching the pattern observed in the sample of homozygous derived individuals, indicating this decline is likely not an artefact of small sample size (Supplementary Figure 3). We identified a recent, post-1980 decline across 3 of the 10 subsamples, which could indicate spatial heterogeneity in recent decline patterns across LG1 homozygous ancestral Northern cod samples (Supplementary Figure 3). To test this hypothesis, we split LG1 homozygous ancestral, LG1 homozygous derived, and heterozygous groups into population groups corresponding to coastal Labrador samples, offshore samples, and samples from coastal Newfoundland. We then conducted LinkNe analysis on these samples, revealing recent decline of LG1 homozygous ancestral and heterozygous individuals within the coastal Labrador grouping (Supplementary Figure 2). Together, these results indicate that small sample sizes have likely not produced false signals of decline, and demonstrate spatial variation in cod stock decline across Newfoundland and Labrador.

### Bottleneck detection

To provide secondary genetic confirmation of observed stock collapses observed in abundance data and using effective size reconstruction, we also tested for presence of recent genetic bottlenecks in LG1 homozygous ancestral, heterozygous, and LG1 homozygous derived groups. We analyzed 6270 genome-wide SNPs excluding LG1 and LG12 rearrangements in each group using the BOTTLENECK^51^ software to detect bottlenecks assuming an infinite alleles model. We inferred bottlenecks from presence of a mode-shift in allele frequency toward reduction in the number of low frequency alleles, or significant sign and standardized differences test results. Bottlenecks were observed using all three methods in homozygous groups (sign test, standardized differences test p < 0.0005), but allele frequency mode shift was not detected in heterozygotes. This observation may be attributed to historical scenario sensitivity of the different tests used in the BOTTLENECK software, or may reflect differences due to recent admixture within heterozygous individuals.

## Acknowledgements

We thank C. Hollenbeck for assistance with LinkNe software. We also thank L. Weir, D. Heath, and R. Waples for helpful comments on the manuscript, and the Centre for Integrative Genetics (CIGENE) for genotyping of Atlantic cod used in this study.

## Author Contributions

TK, SJL, IRB, and PB designed the study. TK, SJL and ES performed statistical analyses. PB, IRB, MSW, RF, MK, and SL provided molecular data for the study. PR provided cod abundance survey data. TK drafted the manuscript and all authors contributed to writing and editing of the final manuscript.

